# Strandings of loggerhead sea turtles south of the Po River delta: insights from a highly impacted area

**DOI:** 10.1101/2022.06.28.497895

**Authors:** Luca Marisaldi, Andrea Torresan, Andrea Ferrari

## Abstract

The northern Adriatic Sea is an important foraging ground for the loggerhead sea turtle *Caretta caretta* (Linnaeus, 1758) within the Mediterranean Sea. Here, spatial-temporal patterns of loggerhead sea turtles strandings along a short portion (∼18 km) of the coast south of the Po River delta (Italy) during a three-year period (2019-2021) were investigated. A total of 244 records (alive, *n*=7; dead, *n*=237) were analysed and the curved carapace lengths (CCL, notch to tip, cm) mainly reflected sub-adults (average CCL=55.2 cm; 95% CI= 53.3-57). The month of July was identified as the critical month with the highest number of strandings, mirroring migratory processes toward this area during warmer months. Interaction with the trawl fishery was hypothesized as the main cause of mortality and a small fraction of deaths (6%; *n*=16) could be linked to boat strikes and net entanglement. The number of stranded turtles•km^-1^ as well as the absolute number of strandings along the short portion of monitored coast confirmed this area as the most impacted in Italy and perhaps in the whole Mediterranean Sea. This study provides valuable information to improve conservation efforts for this species and highlight that, with all due caution, monitoring stranding events can offer useful insights into the geographic ranges, seasonal distribution, and life history of marine species of conservation interest such as the loggerhead sea turtle.

## Introduction

The Adriatic Sea subregion within the Mediterranean Sea is considered an essential foraging ground for the loggerhead sea turtle *Caretta caretta* (Linnaeus, 1758) (Almpanidou *et al*., 2022), the most common sea turtle species present in the Mediterranean sea (Margaritoulis *et al*., 2003). Indeed, in the neritic area of the northern Adriatic Sea (<200 m) both juveniles and adult loggerhead sea turtles, most of them arriving from Greek rookeries, can easily access benthic food resources and they are permanent or at least seasonal residents (Lazar *et al*., 2004; Casale *et al*., 2012; Luschi *et al*., 2013). Furthermore, the western part of the basin (average depth ∼30 m) is a highly productive area that is strongly influenced by the inter-annual freshwater discharges of the Po river, which represent the main source of nutrients in the whole basin and sustains a high biodiversity community that is important for the trophic interactions of sea turtles (Lazar *et al*., 2011; Casale *et al*., 2018). However, the northern Adriatic Sea area is among the most exploited and affected by cumulative impacts on a global scale (Lejeusne *et al*., 2010; Ramírez *et al*., 2018) and it is subject to both direct and indirect human-related threats.

The high density of sea turtles along with intense fishing activities in the area are considered to drive the highest sea turtles by-catch rates in the whole Mediterranean by bottom trawling (Lucchetti & Sala, 2010; Casale, 2011; Lucchetti *et al*., 2016), trammel and gillnets (Lucchetti *et al*., 2017). This is a serious matter of concern due to high mortality rates and gas embolism caused by interactions with fisheries (Tomás *et al*., 2008; Franchini *et al*., 2021). Certainly, other anthropogenic threats also include marine debris and pollution, boat strikes and habitat degradation, which are a matter of concern too for the conservation of this species (Casale *et al*., 2010, 2018). Whilst quantifying and describing the effects of such threats at sea is rather difficult to achieve for logistical constraints, monitoring coastal strandings represents a more feasible approach and a cost-effective solution. Accordingly, a proper monitoring program of stranded sea turtles can provide a fair picture of the situation at sea, delivering useful information about interaction with marine debris (Domènech *et al*., 2019), migration patterns (Tolve *et al*., 2018) as well as delineate areas of high risk (Dimitriadis *et al*., 2022). Therefore, developing a regular monitoring program of strandings is a useful approach that contributes to improving conservation plans. For this reason, we present here the results of a three-year monitoring program (2019-2021) of stranded loggerhead sea turtles along the coast south of the Po River delta, an area within a key foraging ground for this species. Data collection and analysis of stranding events under a monitoring program like the one presented here is a useful approach to inform the policy-making process about the impact of anthropogenic activities on protected marine species and therefore to mitigate threats for better conservation goals.

## Materials and methods

### Data collection

The stranding records presented here were collected for three years (2019-2021) from Lido di Volano (FE) to Lido di Spina (FE), accounting for 23 km of coastline (Fig. 1). Due to operational constraints, the monitored area was resized in July 2020 down to 13 km from Lido delle Nazioni (FE) to Lido di Spina (FE). This aspect was carefully considered during the data analysis to correct for any bias (e.g., calculation of the number of strandings•km^-1^; see next paragraph). According to a well-defined protocol, during beach patrolling or upon notification from citizens and from the local coast guard, sea turtles were located, identified, and visually assessed. For each stranded sea turtle, the following parameters were recorded: (i) species identification (ii) GPS coordinates, (iii) date and time, (iv) if possible, sex, (v) curved carapace length (CCL, notch-to-tip, cm) and width (CCW), (vi) external signs of traumas (e.g., boat collision, net entanglement), infection, or parasites, (vii) if dead, decomposition state, (viii) presence of flipper tags (communication to the organization which applied tags was done upon tag recovery). From June 2020, sea turtles that were found alive and debilitated were recovered at the local first aid centre in Porto Garibaldi (FE), then transferred within 12h to the closest (distance <30 km) rehabilitation centre located in Marina di Ravenna (RA), in compliance with the Italian National Guidelines (ISPRA, 2013). Stranded carcasses in optimal conservation status were delivered to a local public institute (“Istituto Zooprofilattico Sperimentale della Lombardia e dell’Emilia Romagna”) for necropsies.

**Figure 1.**
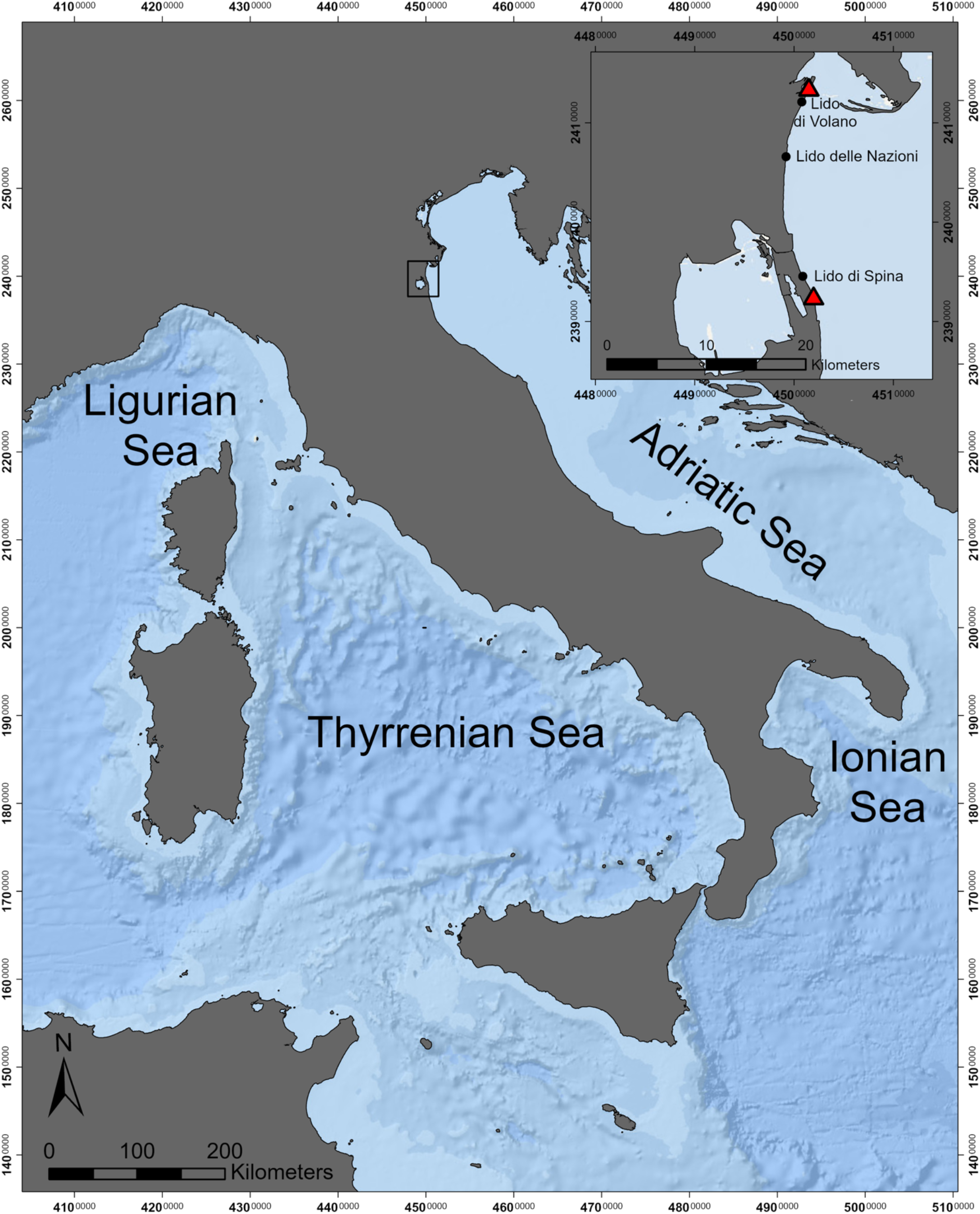
Map showing Italian waters and the coastline as well as the relative position of the study area (black square). The insert (top-right) represents the study area with its extremes (red triangles) and it is provided at higher focus for better details and resolution.

### Data analysis

First, the dataset was explored with a visual approach (i.e., boxplots, scatterplots, Q-Q plot) as a routine step of quality control before applying any statistical analysis. Then, the Shapiro-Wilk and Bartlett’s tests were applied to further check that assumptions for parametric tests were met. Finally, one-way ANOVA and *t*-test were used for comparing CCL among years and between sex, respectively.

The number of strandings•km^-1^ was calculated according to the number of strandings recorded along the kilometres of monitored coast in each month and year. In this regard, due to a resizing during July 2020, the kilometres used to calculate the overall number of strandings•km^-1^ in 2020 were averaged, resulting in 18 km of monitored coast. The statistical and data analysis as well as data visualization was performed within the R environment (R Core Team, 2021). The geographical maps were created with ESRI™ ArcGIS Pro (v. 2.8.0) using the European Terrestrial Reference System 1989, Lambert azimuthal equal-area coordinate reference system.

## Results

### Stranding events and cause of deaths

A total of 255 stranding records were registered from 2019 to 2021 along the coast of the study area, 244 of which could be recorded with CCL and the other parameters included in the protocol. As expected, the most common species was the loggerhead sea turtle *Caretta caretta (n=*254), with only one individual of green sea turtle *Chelonia mydas (*CCL=56 cm). For this reason, all downstream analysis is referred to strandings of loggerhead sea turtles. Flipper tags recovery was low (*n*=2) and their origin was Tunisia (*n*=1) and Italy (*n*=1; Emilia-Romagna). The cause of death could be determined with high confidence only for a small number of cases, which included boat propeller strikes (*n=*15) and net entanglement (*n=1*). The rest of the stranded sea turtles (*n*=228) was absent from any external clear sign of trauma, injury, infection, or parasites.

### Trends

The month with the highest number of records was July for all three years (Fig. 2A), a trend that mirrored the number of stranded turtles•km^-1^ (Fig. 2B). Indeed, during July between 1.3 and 2.6 stranded turtles•km^-1^ were recorded, with the only exception of an isolated peak during November 2021 that reached 1.7 stranded turtles•km^-1^ (Fig. 2B). The months of June, August and September showed lower records than July, although they tended to remain higher than the rest of the year (Fig.1A-B). Interestingly, the overall number of stranded turtles•km^-1^ was 7.4 in 2021, almost twice that of 2019 (3.9) and 2020 (3.8) (Fig. 2C).

**Figure 2.**
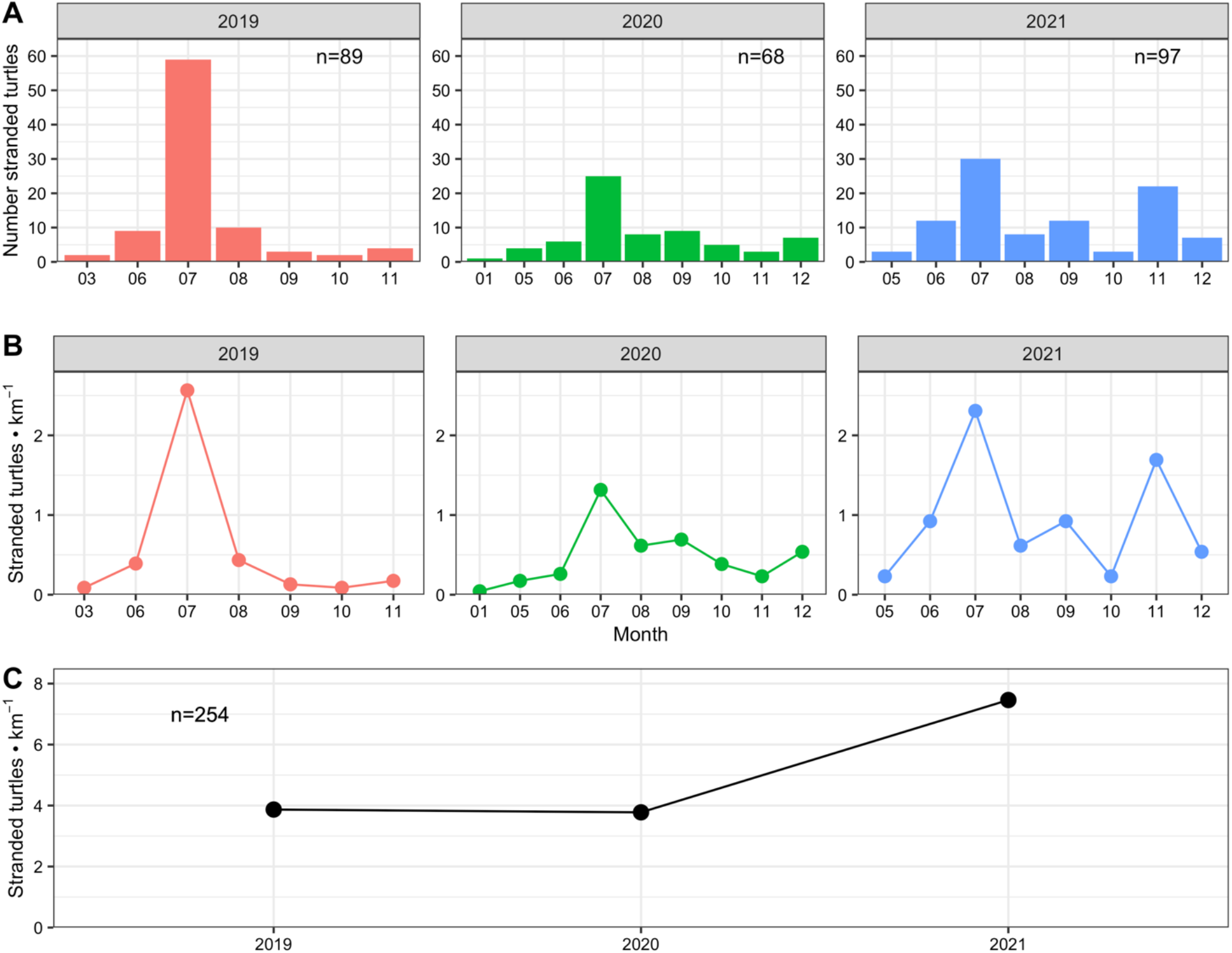
Strandings of loggerhead sea turtles during the three years of monitoring activities in the area (*n*=254). A) Frequency of strandings within each year and month; B) Number of strandings•km^-1^ within each year and month; C) Overall number of strandings•km^-1^ for each year.

The distribution of CCL mainly reflected sub-adults and showed a higher variability during warmer months (June, July, August), with a trend of increasing sizes from September (*n*=12, average CCL=48.1 cm; 95% CI= 42.1-54.4) through December 2021 (*n*=7, average CCL=68.4 cm; 95% CI=57.2-79.6) (Fig. 3A). The CCL was similar among years (one-way ANOVA: F_2,240_=0.1; *p*-value=0.89) and reflected the typical size of sub-adults (Fig. 3B). Accordingly, the average CCL was 55.8 cm (95% CI: 53.1-58.4 cm) during 2019, 54.7 cm (95% CI: 50.6-58.7 cm) during 2020 and 55 cm (95% CI: 51.9-58 cm) during 2021 (Fig. 3B). The number of stranded juveniles (CCL≤30 cm) was low (*n*=7; alive=3, dead=4).

**Figure 3.**
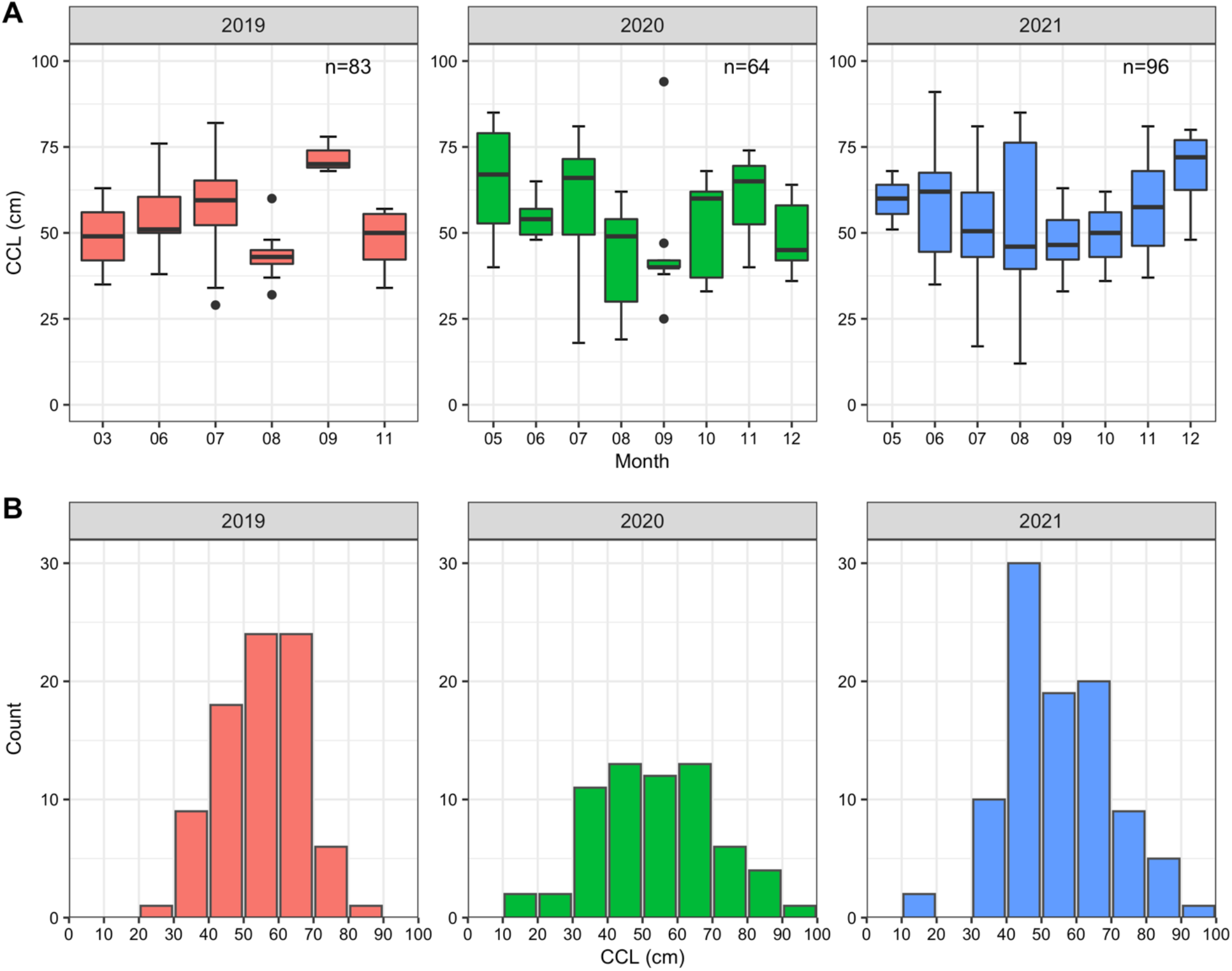
Distribution of CCL (notch to tip, cm) of loggerhead sea turtles during the three years of monitoring activities in the area (*n*=243). A) Boxplots of CCL within each year and month; B) Frequency distribution of CCL in each year.

A total of 32 males and 20 females could be visually identified with high confidence (Fig. 4). The average CCL was 75 cm (95% CI: 71.3-78.7 cm) for males and 73.8 cm (95% CI: 71.6-76 cm) for females, a difference of 1.2 cm (two-sample *t-*test: *t*_*50*_*=-*0.6; *p-*value=0.5).

**Figure 4.**
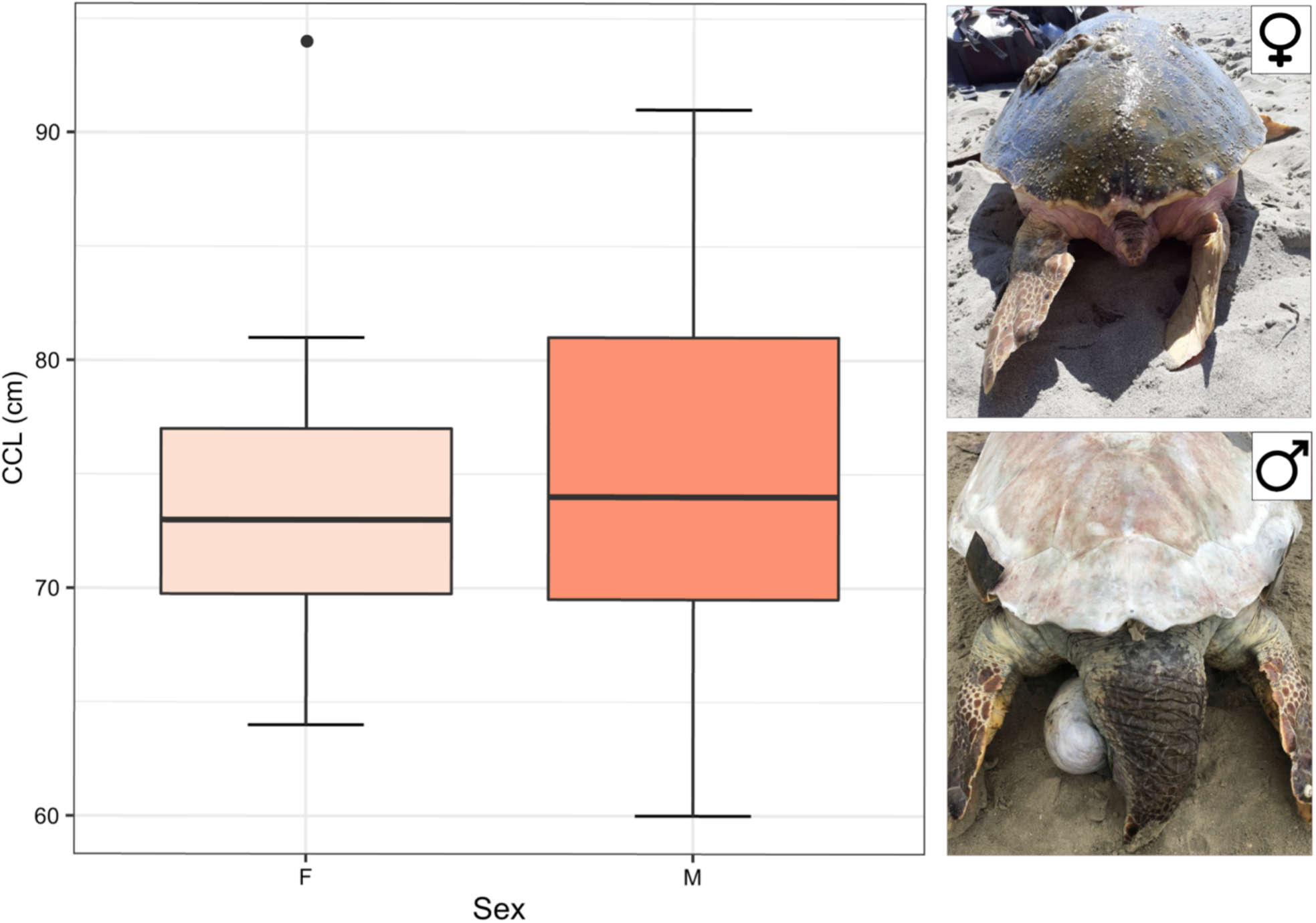
Comparison of CCL (notch to tip, cm) between females (F) and males (M) as well as representative pictures of the tails, one of the main characteristics that allow to distinguish the sex once the adult stage is reached. Photo credit: Andrea Ferrari.

## Discussion

Present results confirm the northern Adriatic Sea as an overwintering area for this species as stranding events of both adults and sub-adults occurred throughout the year, albeit much more frequent during summer months. Sub-adults were recorded more often than adults and the month of July was identified as the most critical month with the highest number of records over the three-year period. Interaction with the trawl (bottom + midwater) fishery was hypothesized as the main cause of mortality. The potential of regular monitoring programs to investigate interactions with direct and indirect anthropogenic stressors (e.g., marine debris, fishing activities) and to detect trends in sea turtle populations is discussed. In the Mediterranean, the sea turtle *Caretta caretta* performs regular migrations from central (i.e., Libya) and central-eastern (i.e., Greece) Mediterranean rookeries to the northern Adriatic Sea (Lazar *et al*., 2004; Bertuccio *et al*., 2019), where shallow waters, transitional habitats and rich benthic communities make it an ideal foraging habitat for this species. Here, both juveniles and adults were found as seasonal or permanent residents, even at low water temperatures (<12 °C) (Lazar *et al*., 2004; Zbinden *et al*., 2008; Casale *et al*., 2012, 2018). Our results further support such findings since stranding events occurred for most of the year, although much more frequent during summer months, a finding in line with a population composed of both residents and migrating individuals. Furthermore, due to its highest number of stranded sea turtles, the month of July might represent the peak of a migratory process toward the area south of the Po River delta, although to date we cannot rule out other drivers for such increased stranding events (i.e., increased fishing effort). Using catch per unit effort (CPUE) of loggerhead sea turtles by the Adriatic midwater pair trawl fishery as an indirect estimate of abundance, Pulcinella et al. (2019) found increased values between April and September in waters near the Po River delta, which suggests migration toward this area during warmer months (Luschi & Casale, 2014). On the other side, autumn was the period with the highest interaction between bottom trawlers and loggerhead sea turtles on the western side of the Adriatic sea (Lucchetti *et al*., 2016). In this context, post-catch direct and potential mortality of trawling (midwater + bottom) in the north Adriatic was estimated at 9.4% and 43.8%, respectively, percentages that likely represented underestimates (Casale *et al*., 2004). Although the exact cause of death can be accurately investigated only with necroscopies, we hypothesise that the majority of strandings were the result of direct interactions with fishing activities (i.e., by-catch in the trawl fishery). Indeed, only a small fraction of deaths could be linked to boat strikes and entanglement in nets, while it was recently found that sea turtles entrapped in static and towed nets may develop gas embolism which can lead to severe organ injury and death, especially if not transported soon at a sea turtle rescue centre (Franchini *et al*., 2021).

From September to December 2021, a trend of stranded sea turtles of increasing size was observed, a result that is consistent with the idea of adult individuals as overwintering residents in the area during that year. However, no trends were noticed during the same months of the previous year, which were instead characterized by a higher presence of sub-adults. It would be interesting to explore in greater detail such year-to-year variability once a more prolonged time series will become available, a step that will allow us to better understand sea turtles strandings in relation to parameters such as sea surface temperature, chlorophyll concentration and fishing effort, an approach recently exploited in the Adriatic Sea for fishery by-caught individuals (Pulcinella *et al*., 2019; Bonanomi *et al*., 2022).

The number of stranded sea turtles•km^-1^ as well as absolute numbers of strandings during three years, clearly confirmed that the area south of the Po River delta is the most impacted in Italy (Casale *et al*., 2010) and perhaps in the whole Mediterranean Sea (Tomás *et al*., 2008; Türkozan *et al*., 2013; Belmahi *et al*., 2020; Hama *et al*., 2020; Dimitriadis *et al*., 2022). Indeed, it is important to highlight that monitoring activities spanned only about 18 km of coastline and the numbers presented herein represented underestimates as we are aware of other and partly overlapping monitoring programs for which data could not be accessed. Furthermore, additional strandings are likely missing in our estimates due to intense beach cleaning activities by a local company during night-time and early morning throughout the summer season, which, to the best of our knowledge, include disposal of sea turtle carcasses. Therefore, we strongly suggest that the current stranding network, data collection and sharing policy in the area must be revised soon to achieve a better understanding of stranding dynamics as well as more precise estimates.

Promising solutions to mitigate such an impact are represented by technological innovations in fishing gear such as the well-known turtle excluder devices (TEDs), already tested with success in the Adriatic sea (Sala *et al*., 2011; Lucchetti *et al*., 2019) but not adopted by fishermen. Eventually, the adoption of TEDs can be coupled with labels that add value to fish products and raise environmental awareness of consumers, who are generally willing to pay a premium price for the attribute of sustainability (Maesano *et al*., 2020). Finally, dynamic spatial-temporal closures is another promising approach that can be more effective at reducing by-catch than classical static closure (Smith *et al*., 2021), although a good understanding of by-catch hotspot must be achieved first (Cambiè *et al*., 2013).

Overall, the results presented here provide valuable information to improve conservation efforts for the loggerhead sea turtle *Caretta caretta* and emphasize a high impact on this species in the area south of the Po River delta, although our hypotheses await confirmations with other approaches. Indeed, albeit inference from strandings must be taken with caution, when investigated over wide temporal scales and considered in conjunction with other approaches, they can offer useful insights into the geographic ranges, seasonal distribution, and life history of marine species of conservation concern such as the loggerhead sea turtle.

## Data availability

The data that support the findings of this study are available from the corresponding author, upon reasonable request.

## Acknowledgements

The authors wish to acknowledge Dr. Alessia Crosara for her valuable support during the creation and visualization of maps. Also, the authors wish to thank Michela Fontana, Noemi Calabretti, Rebecca Montanari and Alessio Alberghini for their precious help during field activities.

